# Maintenance of active chromatin states by Hmgn1 and Hmgn2 is required for stem cell identity

**DOI:** 10.1101/498733

**Authors:** Sylvia Garza-Manero, Abdulmajeed Abdulghani A. Sindi, Gokula Mohan, Ohoud Rehbini, Valentine H. M. Jeantet, Mariarca Bailo, Faeezah Abdul Latif, Maureen P. West, Ross Gurden, Lauren Finlayson, Silvija Svambaryte, Adam G. West, Katherine L. West

## Abstract

Chromatin plasticity is thought to be fundamental to the pluripotency of embryonic stem cells. Hmgn proteins modulate chromatin structure and are highly expressed during early development and in neural stem/progenitor cells of the developing and adult brain. Here, we show that loss of *Hmgn1* or *Hmgn2* in pluripotent embryonal carcinoma cells leads to increased levels of spontaneous neuronal differentiation. This is accompanied by the loss of pluripotency markers and increased expression of the pro-neural transcription factors *Neurog1* and *Ascl1*. Neural stem cells derived from these *Hmgn-*knockout lines also show increased spontaneous neuronal differentiation and *Neurog1* expression. The loss of Hmgn2 is associated with the disruption of active chromatin states at specific classes of gene. The levels of H3K4me3, H3K9ac, H3K27ac and H3K122ac are considerably reduced at the pluripotency genes *Nanog* and *Oct4*, which impacts transcription. At endodermal/mesodermal lineage-specific genes, the loss of *Hmgn2* leads to a switch from a bivalent to a repressive chromatin configuration. However, at the neuronal lineage genes *Neurog1* and *Ascl1*, no epigenetic changes are observed and their bivalent states are retained. We conclude that Hmgn proteins play important roles in maintaining chromatin plasticity in stem cells, and are essential for maintaining stem cell identity and pluripotency.

## Introduction

The chromatin landscape of embryonic stem cells has distinct characteristics that are believed to confer increased plasticity and thus contribute to the maintenance of the pluripotent state (1). Plasticity is achieved through a range of mechanisms, including bivalent epigenetic states (2), more numerous DNaseI hypersensitive sites (3), increased levels of active transcriptional marks, fewer and less condensed heterochromatin loci, and hyperdynamic binding of major architectural proteins such as the linker histones (4).

Hmgn (High Mobility Group Nucleosome binding) family members are chromatin architectural proteins that influence local and global chromatin structure, and have wide ranging effects on gene expression (5). They are devoid of enzymatic activity, and bind to the nucleosome or core particle of chromatin without specificity for the underlying DNA sequence. The two most abundant and well-studied Hmgn isoforms are Hmgn1 and Hmgn2. The five Hmgn family members share a general structure, consisting of a conserved positively charged nucleosome binding domain (NBD) that specifically targets the nucleosome and constitutes the hallmark of the family, a bipartite nuclear localisation signal, and a variable negatively charged C-terminal or regulatory domain (RD) (6).

Hmgn proteins have been shown to modulate chromatin structure in a variety of ways. They can alter the ability of histone acetylases and kinases to modify core histone N-terminal tails (7–9) and have been shown to inhibit the activity of chromatin remodelling complexes (10). They also inhibit the interaction of linker histones with chromatin, which can lead to decompaction and reduced EZH2-mediated methylation of histone H3K27 (11,12). They have also been shown to increase the recruitment of certain transcription factors to chromatin (13).

Chromatin immunoprecipitation studies from several labs have shown that Hmgn proteins do not tend to be highly enriched at individual genes (9,14–16). This is consistent with observations that Hmgns do not have any specificity for the underlying DNA sequence, and that their binding to chromatin is transient and dynamic, and with continuous exchange from one nucleosome to another (17). However, ChIP-seq studies by the Bustin lab have shown that Hmgn1 and Hmgn2 are enriched at DNase I hypersensitivity sites (DHSs) in several mammalian cell lines (3,18–20). DHSs constitute chromatin accessible domains located at transcriptional regulatory regions, such as promoters and enhancers, and are considered to be a hallmark of genes poised or activated for transcription. Cells lacking both Hmgn1 and Hmgn2 exhibit a reduction in the number and the intensity of the DHSs in mouse embryonic fibroblasts and mouse embryonic stem cells (ESCs) (19,20).

The generation of *Hmgn* variant-specific knockout mice in the last decade has provided considerable insights into the relevance of these proteins at the cellular and organism level. In general, these mice are viable and do not present strong phenotypes. Nevertheless, variant-specific phenotypic alterations have been reported. For example, *Hmgn1*^−/−^ mice exhibit increased tumorigenicity and impaired DNA damage response (21), whereas *Hmgn2*^−^*^/^*^−^ mice show some alterations in energy metabolism (19). Analyses of transcriptional changes in cells and tissues derived from the genetically modified mice indicate that the absence of one or two *Hmgn* variants does not dramatically modify the pre-existing transcriptional profile (3,12,19,20). However, there are significant changes in the levels of many mRNA transcripts, which indicates that the general process of transcription is altered, and accounts for the *Hmgn* knockout mouse phenotypes mentioned above. These experiments led to the working hypothesis that the Hmgns fine tune an already established expression profile, and ensure the appropriate cellular response to external and internal cues required for proper organism functioning.

During development, Hmgn1 and Hmgn2 are highly abundant in embryonic tissues, and then are progressively down regulated as differentiation proceeds, remaining at lower levels in fully differentiated tissues (22). This decreasing expression pattern has been observed during erythropoiesis, myogenesis, and chondrogenesis, and seems to be essential, since Hmgn1 overexpression inhibits normal cellular differentiation (15,23,24). Although fully differentiated cells have lost substantial amounts of Hmgn proteins, many tissue-specific stem cells and transient amplifying precursors retain high levels of these proteins (22,25).

In the mouse brain, *Hmgn1* and *Hmgn2* mRNAs are highly expressed in the dentate gyrus of the adult hippocampus (3), which is a well characterised neurogenic niche where neural stem cells (NSCs) reside and undergo active neurogenesis. A role for Hmgn proteins in NSCs is supported by the observation that adult *Hmgn1*^−^*^/^*^−^ mice exhibit a reduction in Nestin-positive cells in the sub-ventricular zone (3). Moreover, Hmgns are required for proper differentiation of NS cells into astrocytes and oligodendrocytes, since Hmgn knockdown interferes with the neuron to glia transition, and the lack of both Hmgn1 and Hmgn2 leads to down-regulation of Olig1 and Olig2 and defects in oligogenesis (12,26). Defects in neural stem cell differentiation could be a contributing factor to the neurological defects observed in *Hmgn1*^−^*^/^*^−^ and double knockout *Hmgn1*^−^*^/^*^−^*Hmgn2*^−^*^/^*^−^ mice (12,27). Together these data suggest a role for Hmgn proteins in tissue-specific stem cell biology and differentiation, especially among the neural lineage.

In this study, we have investigated the role of Hmgn1 and Hmgn2 in P19 embryonal carcinoma stem cells. P19 cells are pluripotent embryonal carcinoma cells that maintain their phenotype when cultured in the presence of serum. They were originally derived from post-implantation embryos (28), capturing a later developmental stage than mouse ES cells. P19 cells resemble epiblast stem cells, a primed state of pluripotency which shares several features with human ES cells (29–31). P19 cells can be easily differentiated down the neuronal or cardiomyocyte lineages, and have been extensively studied over the last three decades. Our data indicate that Hmgn proteins are important for regulating the active epigenetic landscape and gene expression profile, and are required for stem cell self-renewal and the prevention of inappropropriate differentiation.

## Materials and Methods

### Cell culture

Mouse embryonic carcinoma cell line, P19, was maintained in advanced DMEM/F12 containing 10% NBCS and 1X GlutaMAX. Adherent neural induction was performed using the method of Nakayama *et al*. (32), which involved seeding cell onto laminin-coated culture dishes in neural induction media: DMEM/F12 containing sodium pyruvate and NEAA supplemented with 1 X N2 supplement, 200 mM L-glutamine, 500 nM retinoic acid, 10 ng/ml FGF8 and 10 μM DAPT.

To derive neural stem cells, the adherent neural induction protocol was initiated, and after three days, cells were switched to gelatine-coated flasks in N2B27 medium containing 10 ng/ml FGF2 and 10 ng/ml EGF (33). Cells were passaged eight times before being assayed for NSC characteristics. To differentiate the parental NSCs, cells were plated on laminin-coated dishes in N2B27 without FGF and EGF, and assayed after four days.

### Signalling inhibition

Cells were seeded at a density of 2 × 10^5^ cells/well in 6 well plates, allowed to attach overnight and then 10 μM inhibitor was added: XAV-939 for WNT inhibition (S1180-SEL Stratech), DAPT for Notch inhibition (D5942 Sigma-Aldrich), and SU-5402 for FGFR inhibition (SML0443 Sigma-Aldrich). RNA was collected after 24 hours. For immunofluorescence, cells were plated in the presence of WNT inhibitor and assayed 48 hours later.

### CRISPR mutagenesis

A dual CRISPR nickase strategy (34) was used to generate frameshift mutations in exon I of *Hmgn2* (supplementary figure S1). A plasmid expressing the nickase mutant Cas9D10A-T2A-EGFP was transfected alongside plasmids expressing two gRNAs targeting exon I and intron I of *Hmgn2* (gRNA sequences listed in supplementary table 1). Clonal lines were screened for Hmgn2 protein expression by immunofluorescence and western blotting. PCR followed by TA cloning and sequencing was used to confirm mutagenesis of both alleles in each clonal line. Lines N2-a, N2-b and N2-c carry different genetic changes, and so are independent of each other. To generate frameshift mutations in exon I of *Hmgn1*, a plasmid expressing wild type Cas9-T2A-EGFP was co-transfected alongside two gRNA expression plasmids, one expressing targeting *Hmgn1* and the other targeting *Hypoxanthine Phosphoribosyltransferase 1 (Hprt)*. Selection with 10 μg/ml 6-thioguanine was used to isolate clonal lines with inactivating *Hprt* mutations, in order to enrich for cells in which active CRISPR-mediated mutagenesis had occurred. Lines lacking Hmgn1 expression were identified as above. Lines N1-a and N1-b were independently derived from separate transfections.

### Gene expression analysis

cDNA synthesised from purified RNA using Superscript III (Life Technologies), and qRT-PCR was performed using SYBR green Fast start universal master mix (Roche). The housekeeping gene *Gpi1* was used as the normaliser in ΔΔCt calculations. qRT-PCR primers for *Hmgn1* span the exon V-VI junction, and primers for *Hmgn2* span the exon IV-V junction. Primer sequences are listed in supplementary table 1.

### Western blotting

The Hmgn1 and Hmgn2 antibodies were raised in rabbit against the C-terminal peptides NQSPASEEEKEAKSD and KTDQAQKAEGAGDAK, respectively, and affinity purified (Eurogentec). For western blotting, whole cell lysates were run on 15% SDS-PAGE gels, blotted onto PVDF and membranes probed with the relevant antibodies.

### Immunofluorescence

Cells were fixed with 4% PFA for 30 mins, permeabilised for 10 min with 0.1% PBS-triton X-100, washed with 20 mM PBS-glycine and blocked with 5% horse serum in 0.5% PBS-triton X-100. Primary antibodies from Abcam were against Oct4 (Ab19857), Nanog (Ab80892), Ssea1 (Ab16285), Gata4 (Ab84593), Nestin (Ab6142), βIII tubulin (Ab18207). Secondary antibodies were from Thermo Fisher Scientific (A21206, A11037, A11020). Images shown are representative of the whole slide.

### Chromatin immunoprecipitation (ChIP)

Adherent cells were cross-linked in DMEM/F12 without serum for 5 min with 0.5% formaldehyde. ChIP was performed as described in (20). Briefly, for each chromatin immunoprecipitation (IP) reaction, 5 μg of chromatin and 50 μl of magnetic DYNA beads-Protein A (10002D Thermo Fisher Scientific) were used. Magnetic beads were blocked with 0.5% BSA prior to incubating with 2–3 μl of antibody in 200 μl of blocking buffer. Antibodies were from Millipore: H3K4me3 (07–433), H3K27me3 (07–449), H3 CT pan (07–690), H3K27ac (07–360), H3K4me1 (07–436) or Abcam: H3122ac (ab33309). qPCR reactions contained 2% of IP DNA or 0.5 ng of input DNA. Primer sequences and locations relative to each transcription start site (TSS) are listed in supplementary table 1. The ratio of the PCR readings in IP vs. input DNAs (ΔCt) were calculated and converted to fold enrichments. The average fold enrichment for H3 across all primer sets tested was used to normalise the data for all ChIP reactions, which controls for small differences in chromatin concentration between samples.

For ChIP-seq, libraries were prepared with the NEB ultra DNA library prep kit and single-end sequencing to 76 bp read length was carried out by the University of Glasgow Polyomics facility. Unfiltered reads from FASTQ files were aligned to reference genome NCBI build 37 mm9 using Bowtie (35); reads aligning to more than three positions and duplicate reads were discarded. Statistically enriched regions compared to a background control (peak finding) were identified using MACS (36) with parameters p = 1e-3; bandwidth = 200; genome = mm9; fold enrichment = 10,30; and input control was used as a background.

## Results

### Generation of Hmgn2 and Hmgn2 knockout cell lines

CRISPR mutagenesis (34) was used to create clonal derivatives of P19 embryonal carcinoma cells in which Hmgn1 or Hmgn2 protein expression was abolished (supplementary figure S1). Two independent lines show complete loss of Hmgn1 expression: N1-a and N1-b (Figure 1a). Control line CON-b went through the same process but *Hmgn1* was not targeted and no loss of Hmgn1 protein is observed. Three independent lines show complete loss of protein expression: N2-a, N2-b and N2-c (Figure 1b). Control line CON-a was generated at the same time but has no mutagenesis of the *Hmgn2* alleles.

**Figure 1.**
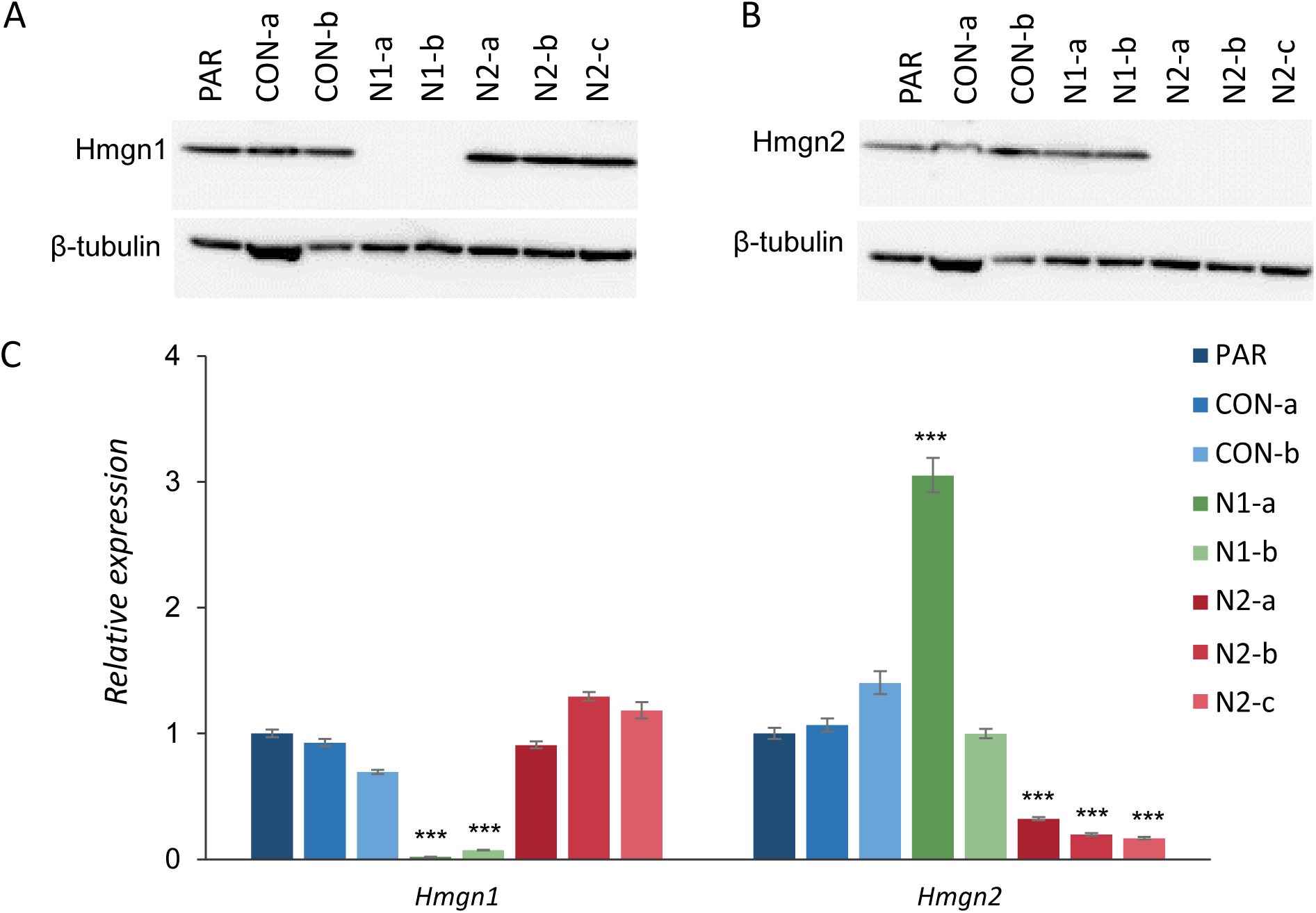
Generation of *Hmgn1* and *Hmgn2* knockout P19 lines. Western blot analysis of (A) Hmgn1 and (B) Hmgn2 protein levels in clonal P19 lines; β-tubulin is shown as a loading control. (C) *Hmgn1* and *Hmgn2* mRNA expression as determined by qRT-PCR in control and Hmgn-knockout cells. The graph represents the average fold change relative to parental P19 cells. Error bars represent the SEM from 3–10 independent cultures. The statistical significance was calculated by ANOVA and Dunnett’s multiple comparison test (adjusted p value ***<0.0001).

Immunofluorescence confirmed the loss of Hmgn1 and Hmgn2 protein in all cells in their respective knockout lines (supplementary figure S2). Quantitative RT-PCR revealed that *Hmgn1* and *Hmgn2* transcript levels are significantly reduced in the knock-out cell lines, indicative of either a change in splicing preferences and/or a reduction in transcript stability (Figure 1c). The level of the *Hmgn2* transcript is increased by three fold in the *Hmgn1* mutant line N1a (Figure 1c), but no corresponding increase in Hmgn2 protein is observed (Figure 1b). No other reciprocal changes in Hmgn1 or Hmgn2 expression were observed in the other knockout or control lines (Figure 1a-c).

### Hmgn knockout cells have reduced expression of pluripotency markers

P19 cells have lost the capacity of spontaneously differentiate, despite their teratocarcinoma origin, and can be propagated indefinitely in serum-containing medium as mostly pure cultures of undifferentiated cells. Parental P19 cells are morphologically homogeneous and grow as discrete colonies. We observed that the loss of Hmgn1 or Hmgn2 expression considerably alters the cellular morphology and organisation of P19 cultures (supplementary figure S3). Fewer discrete colonies are observed, with a high proportion of *Hmgn*-knockout cells growing outside of normal colony boundaries. Many of these cells exhibit an extended cytoplasm and resemble differentiated cells, such as endothelial flat cells (37). Some cells have long processes that connect with other cells (supplementary figure S3).

We hypothesised that these changes in cell morphology in the *Hmgn* knockout populations could reflect an increased propensity to differentiate. In order to investigate this further, the expression of self-renewal markers was assayed by immunofluorescence and qRT-PCR. The core pluripotency transcription factor *Nanog* is highly expressed in naïve pluripotent cells and is down-regulated upon lineage-priming (29,30). In the *Hmgn* knockout lines N1-b, N2-a and N2-c, *Nanog* mRNA expression is reduced by 3–10 fold relative to parental P19 cells (Figure 2a). By immunofluorescence, it can be seen that Nanog protein is heterogeneously expressed by parental and CON-a cells, with brighter cells (arrows) and paler cells (arrow heads) present in the same colony (Figure 2c). This is in agreement with previous reports suggesting that heterogeneity in Nanog production is related to diverse differentiation potentials among a polypopulation of pluripotent cells (38). Notably, Nanog protein expression is reduced in all the *Hmgn1* and *Hmgn2*-knockout cultures (Figure 2c). Most of the *Hmgn1* and *Hmgn2* knockout cells have little or no Nanog expression, although a minor subset of cells retain modest Nanog expression. A reduction in *Nanog* protein in N2-b and N2-c cell lines was also observed by western blotting (Figure 2b).

**Figure 2.**
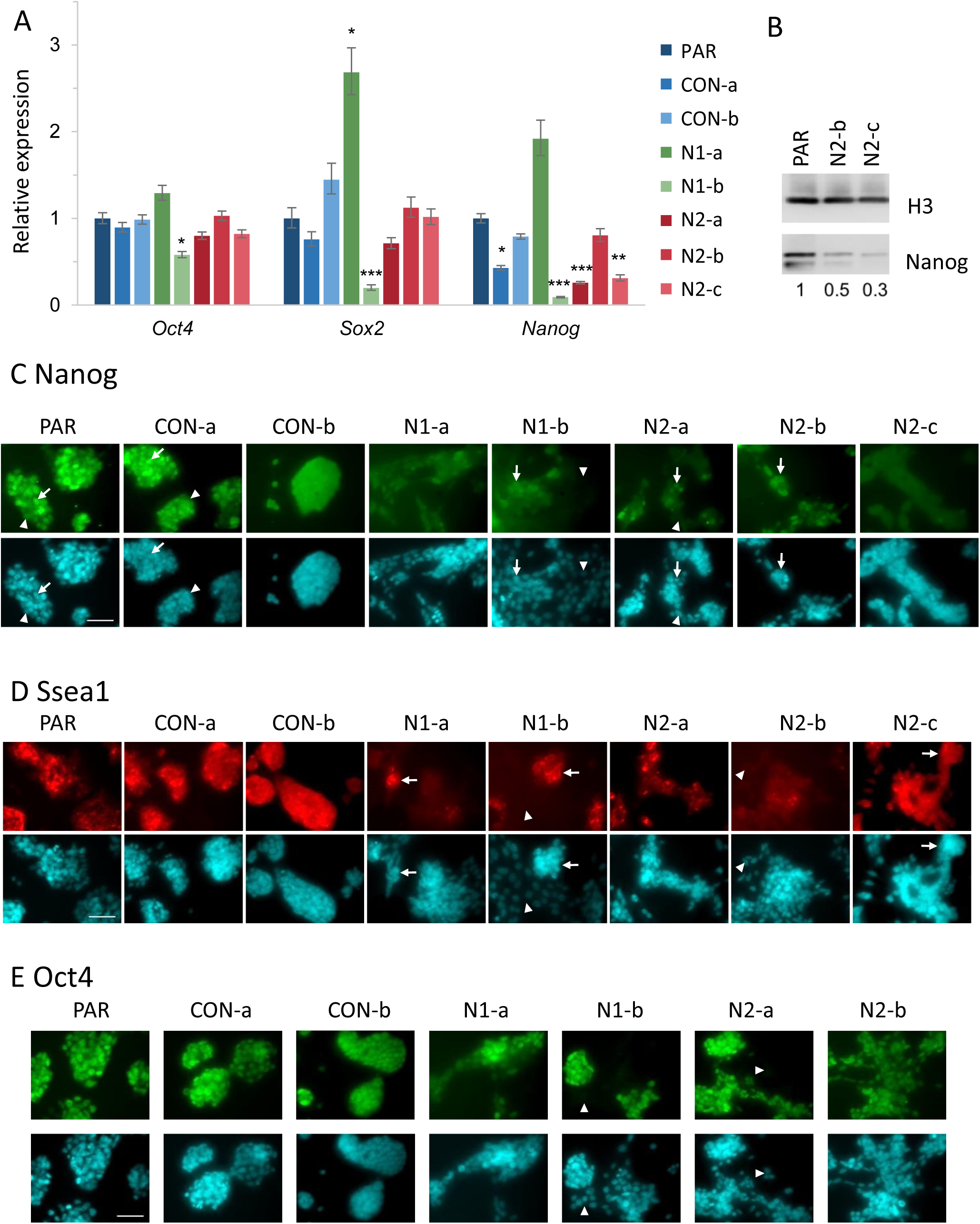
Expression of pluripotency markers in *Hmgn* knockout cells. (A) Relative expression of the pluripotency transcription factors *Oct4, Sox2*, and *Nanog* as determined by qRT-PCR. The graphs show the fold change relative to parental P19 cells. Error bars represent the SEM from 3–10 independent cultures. The statistical significance was calculated by ANOVA and Dunnett’s multiple comparison test (adjusted p values *<0.05, **<0.01, ***<0.001). (B) Western blot analysis of Nanog protein levels in whole cell extracts from parental P19, N2-b and N2-c cells. H3 is shown as a loading control. The ratio of Nanog to H3, relative to that in parental cells, is shown below. (C-E) Immunofluorescence analysis of pluripotency marker expression. Parental, CON-a, and Hmgn-knockout cells were fixed 24 h after seeding. DAPI was used to stain the nuclei (cyan). Scale bar indicates 50 μm. (C) Nanog immunostaining in green. Nanog levels are heterogeneous in all cell lines, as marked by higher (arrows) and reduced (arrow heads) fluorescence intensity. (D) Ssea1 immunostaining in red. Clustered Hmgn-knockout cells retain higher levels of Ssea1 (arrows) than the cells that grow spread outside the colonies (arrow heads). (E) Oct4 immunostaining in green. Arrowheads indicate cells with reduced Oct4 staining.

Stage-specific embryonic antigen 1 (Ssea1) plays a role in cell adhesion and migration in the pre-implantation embryo, is widely used as a pluripotency marker in mouse ESCs, and is also expressed in embryonal carcinoma cells, including P19 cells (Figure 2d) (39). The majority of *Hmgn*-knockout cells lose expression of this pluripotency marker, especially those that grow beyond normal colony boundaries (arrow heads, Figure 2d). A subset of *Hmgn*-knockout cells growing in clusters retain modest Ssea1 expression (arrows), which is lower than that seen in parental, CON-a and CON-b cultures.

*Oct4* is a transcription factor that is highly expressed in undifferentiated pluripotent stem cells. It is has also been shown to be retained in the first stages of ESC differentiation (40). *Oct4* mRNA levels are similar across all the Hmgn knockout lines except N1-b, in which it is decreased by 40% (Figure 2a). This is consistent with immunofluorescence data (Figure 2e). In the parental cells, *Oct4* is homogeneously distributed along all cell lines and most of the cells show equivalent fluorescence intensity. In most of the Hmgn knockout cells, *Oct4* expression in unchanged. However, some cells growing beyond colony boundaries are completely devoid of *Oct4*, especially in N1-b cultures (Figure 2e, arrow heads).

*Sox2* has a similar expression pattern to *Oct4* in the different pluripotency states and during early ESC differentiation (40), and is also expressed in neural stem cells. *Sox2* mRNA expression is not altered in the *Hmgn2* knockout lines, although its expression is increased in N1-a cells and decreased in N1-b cells (Figure 2a).

Taken together, the reduction in Ssea1 and Nanog protein levels suggest that the maintenance of self-renewal and pluripotency are compromised in Hmgn-knockout cultures, even though the cell lines can still be propagated indefinitely.

### Hmgn knockout cells show increased spontaneous neuronal differentiation

To investigate whether *Hmgn* knockout cells show increased levels of spontaneous differentiation in undifferentiated culture conditions, the expression of neuronal, endodermal and mesoderm markers was examined. βIII tubulin immunostaining was performed to detect differentiation down the neuronal lineage (Figure 3a). Strikingly, 24 hours after seeding the cells at low density, all five Hmgn knockout lines showed increased formation of βIII tubulin-positive cells with extended processes, a phenotype that is typical of immature neuronal cells. This contrast strongly with the low frequency of βIII tubulin-positive cells that are typically observed in the control cultures (Figure 3a). At 48 hours after seeding, extensive networks of neurites are visible in lines N1-a, N2-a, N2-b and N2-c (Figure 3a). *βIII tubulin* mRNA levels show an increase of 2 – 2.6 fold in N1-a, N2-b and N2-c cells (Figure 3b)

**Figure 3.**
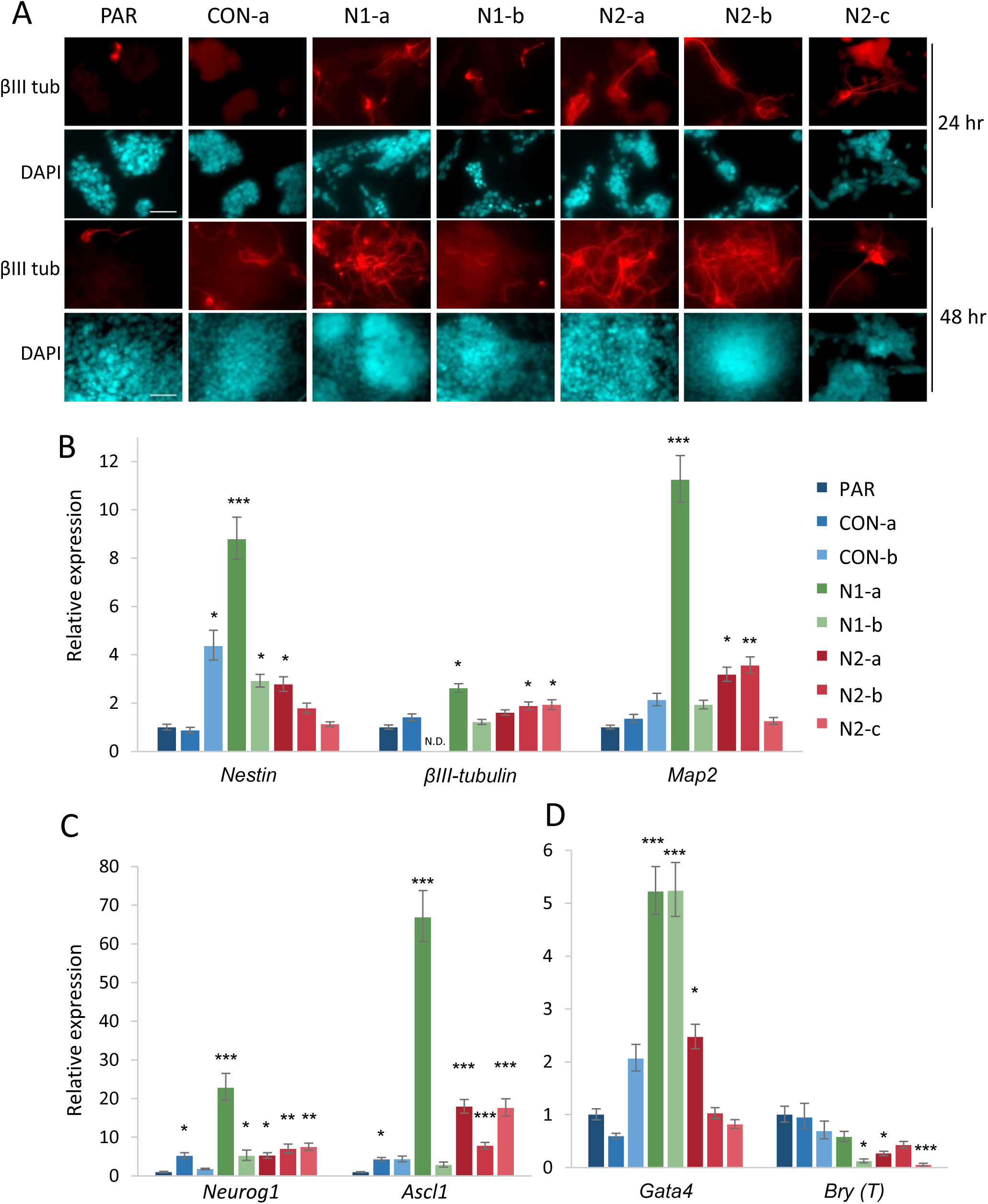
*Hmgn* knockout cells show increased levels of spontaneous neuronal differentiation. (A) Immunofluorescence analysis of the neurofilament protein βIII-Tubulin (red). Parental, CON-a, and Hmgn-knockout cells were fixed 24hr and 48 hr after seeding. DAPI was used to stain the nuclei (cyan). Scale bar indicates 50 μm. (B) Relative mRNA expression of the neural-specific markers *Nestin, βIII-tubulin* and *Map2*, and (C) the lineage-specific transcription factors *Neurog1, Ascl1*, and (D) the endodermal/mesodermal markers *Gata4*, and *Bry* (*T*). N.D: no data available. The graphs show the fold change relative to parental cells. Error bars represent the SEM from 3–10 independent cultures. The statistical significance was calculated by ANOVA and Dunnett’s multiple comparison test (adjusted p values *<0.05, **<0.01, ***<0.001).

Nestin is an intermediate filament that is as a classical marker of neural stem cells. P19 undifferentiated cells express Nestin at basal levels (41). *Nestin* mRNA expression is increased by 3 to 9 fold in N1-a, N1-b and N2-a cells (Figure 3b). Consistent with this, immunofluorescence shows that Nestin protein levels are somewhat higher in the *Hmgn* knockout cultures than in the control lines, particularly in the cells growing in three dimensional clusters (supplementary figure S4, part a, arrows).

*Neurog1* and *Ascl1* are pro-neural transcription factors that play important roles in driving neuronal differentiation (42). Consistent with the increased spontaneous neuronal differentiation of Hmgn-knockout cells, *Neurog1* and *Ascl1* are upregulated in most of the *Hmgn*-knockout lines when compared with parental cells (Figure 3c). *Neurog1* transcription is increased by 5 to 25 fold in *Hmgn1*-knockout and *Hmgn2*-knockout lines, and *Ascl1* is increased by 8 to 18 fold in *Hmgn2-* knockout cells and 67 fold in N1-a cells when compared with the parental cells (Figure 3b).

P19 cells can be differentiated down the mesodermal lineage into cardiomyocytes in the presence of DMSO or other inducers (43). To test whether *Hmgn*-knockout cells spontaneously differentiate down this lineage, *Gata4* and *Brachyury* (*Bry or T*) expression was analysed. *Gata4* and *Brachyury* are typically regarded as mesodermal and/or endodermal markers. In differentiating P19 cells, *Brachyury* is an early marker of mesodermal differentiation, whereas *Gata4* is expressed slightly later, in cardiomyoblasts, where it regulates cardiomyocyte differentiation (44). We find that *Gata4* transcript levels are increased by around 5 fold in the *Hmgn1* knockout lines N1-a and N1-b, but are only increased by 2.5 fold in one of the *Hmgn2* knockout lines, N2-a (Figure 3d).

Immunofluorescence reveals very few Gata4-positive cells are across all the control and Hmgn knockout cell lines (supplementary figure S4b). However, a subset of Gata4-positive cells are observed in line N1-b (Figure S4b, arrows). Conversely, *Brachyury* transcript levels are decreased in three of the Hmgn knockout cell lines (N1-b, N2-a and N2-c) (Figure 3d). Although the increase in *Gata4* expression might indicate initiation of cardiogenesis, this is not consistent with the reduction in *Brachyury* expression. Therefore, this data may be more consistent with a general loss of control of self-renewal in the *Hmgn1* knockout cells.

Taken together, this data shows that the loss of either *Hmgn1* or *Hmgn2* compromises the self-renewal of pluripotent embryonal carcinoma cells, disrupts the gene expression profile, and leads to the initiation of differentiation programs that are mainly, but not exclusively, towards neural lineages.

### Spontaneous neuronal differentiation is not driven by increased autocrine signalling

The main signalling pathways involved in neuronal differentiation are Notch signalling, which promotes survival and self-renewal of NS cells (45), FGF signalling, which is important for both NS cell proliferation and differentiation (46), and WNT signalling, which can either promote self-renewal or neuronal specification, depending on the developmental stage (47). We hypothesised that these pathways may be dysregulated in the *Hmgn* knockout cells, leading to the increased levels of spontaneous neuronal differentiation that were observed.

To test this hypothesis, three known targets of these pathways were studied, *Axin2, Fgf4* and *Hes5*. Activation of WNT signalling promotes a negative regulatory feedback loop, in which the expression of the canonical WNT inhibitor *Axin2* is up-regulated (47). FGF signalling activates transcription of *Fgf4*, which promotes neuronal differentiation and is detected in the early stages of P19 neuronal differentiation (46). Hes5 is a basic helix loop helix transcription factor whose transcription is activated by Notch signalling. It represses pro-neuronal transcription factor gene expression and thus prevents the differentiation of NS cells (48).

The expression of these target genes in the *Hmgn2* knockout lines was evaluated by qRT-PCR (Figure 4a). No significant changes in expression of *Hes5, Axin2* or *Fgf4* were observed in the *Hmgn2* knockout cell lines, indicating that the activity of these signalling pathways is not altered. To confirm that these pathways are active in the P19 lines, cells were treated with the Notch signalling inhibitor DAPT, the FGF receptor inhibitor Su05402, or the WNT pathway inhibitor XAV-939. In both control and *Hmgn2* knockout cell lines, the signalling inhibitors reduced expression of the effector genes *Axin2, Fgf4* and *Hes5* as expected (Figure 4a).

Although Notch, FGF and WNT signalling does not appear to be increased in *Hmgn2* knockout cells relative to the parental cells, it is possible that the knockout cells respond differently to the signalling events that are active. For example, changes in chromatin structure could mean that target genes are more or less responsive to a given level of signalling. Therefore, we investigated the effects of inhibiting these signalling pathways on gene expression and cellular phenotype. As *Nanog* expression is reduced in several of the *Hmgn* knockout lines, we investigated whether its expression is altered following treatment with signalling inhibitors. It can be seen that the profile of changes in *Nanog* expression is the same in the *Hmgn2* knockout lines as it is in the control lines (Figure 4b). Specifically, *Nanog* expression is reduced following treatment with the WNT inhibitor, unchanged following Notch inhibitor treatment, and increased after treatment with the FGF inhibitor. We previously showed that *Neurog1* expression is increased in *Hmgn2* knockout lines (Figure 3c). Treatment with signalling inhibitors has similar effects in *Neurog1* expression in both the knockout and the control lines: *Neurog1* expression is unchanged when WNT signalling is inhibited, slightly increased (up to 3 fold) when Notch signalling is inhibited, and increased by 7 – 13 fold when FGF signalling is inhibited (Figure 4b).

Finally, immunofluorescence was performed to investigate whether the proportion of neuronal cells was altered following inhibition of WNT signalling. It can be seen that the WNT signalling inhibitor does not substantially reduce the number of neuronal cells in N2-b and N2-c populations (Figure 4c). Taken together, this data indicates that the autocrine WNT, FGF and Notch signalling pathways are not enhanced in *Hmgn2* knockout cells, and that the increased spontaneous neuronal differentiation does not appear to be driven by these signalling pathways.

### Induced neuronal differentiation proceeds normally in *Hmgn1* and *Hmgn2*-knockout cells

In order to investigate whether Hmgn knockout cells show an increased ability to differentiate into neurons following induction stimuli, a well-established P19 neuronal induction protocol was used (32). This includes the stimulation of the cells with retinoic acid and fibroblast growth factor 8 (FGF-8), and the use of the Notch pathway inhibitor DAPT to stimulate cell cycle exit. Expression of neuronal lineage genes including *Nestin, Neurog1* and *Ascl1* is strongly induced by day 3, and complex networks of neurites are apparent by day 4 (Figure 5a and b, supplementary figure S5) (32).

**Figure 4:**
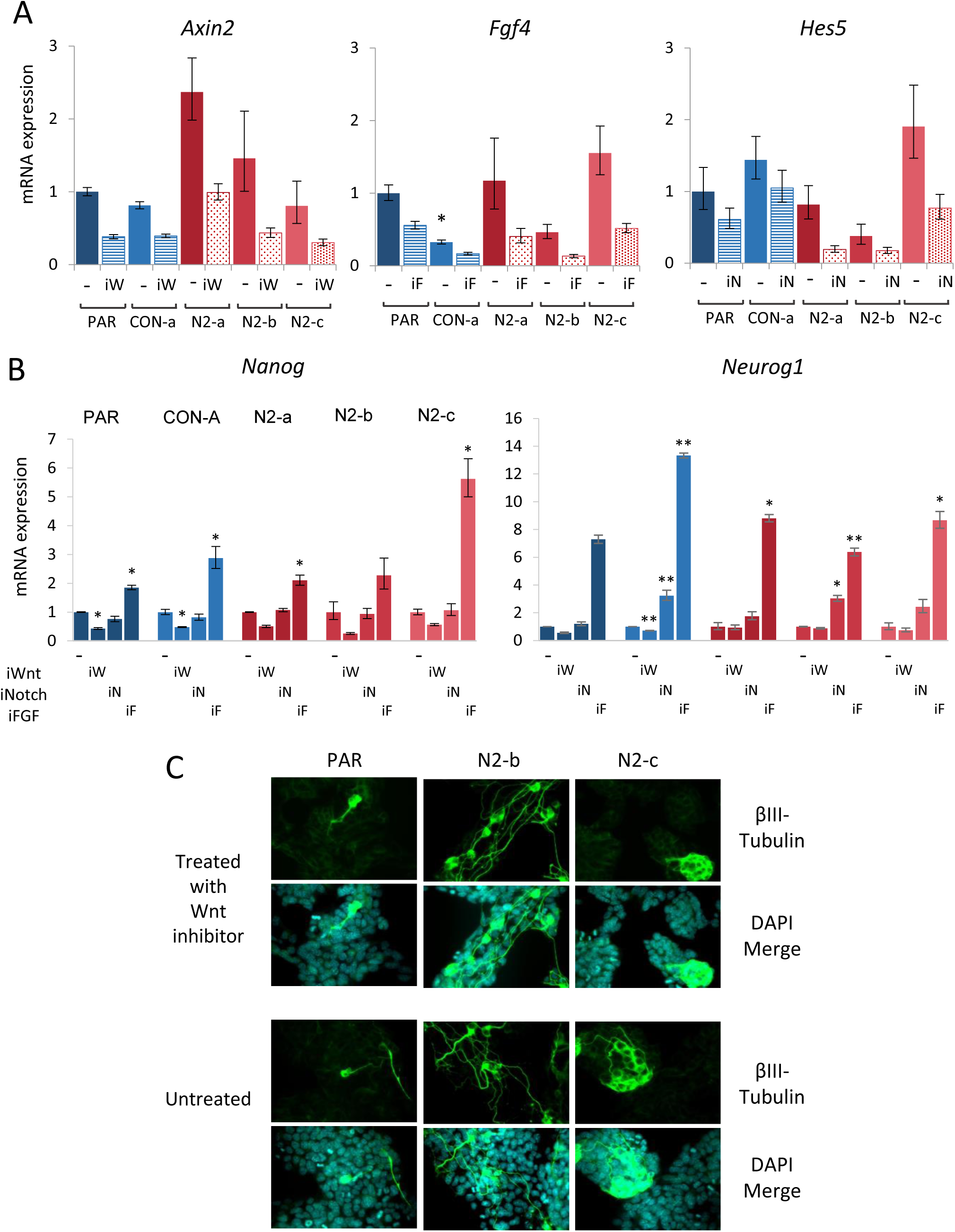
Inhibition of WNT, Notch or FGF signalling pathways does not alter the *Hmgn2* knockout cell phenotype. (A) Relative expression of the reporter genes for WNT (*Axin2*), Notch (*Hes5*) and FGF (*Fgf-4*) signalling pathway activity and (B) relative expression of *Nanog* and *Neurog1* in parental, CON-a, and *Hmgn2-*KO cells, as determined by real time RT-PCR. Expression levels of the reporter genes were assayed after 24 h of treatment with 10 μM of the signalling pathway inhibitor: DAPT for Notch, SU-5401 for FGFR, and XAV-939 for WNT. The graphs present the mean and s.d. of 3 independent cell cultures per cell line. In (A), expression is plotted relative to that in untreated parental cells. The Student’s T-test was used to test for significant differences between untreated parental and untreated control or knockout lines. * = P < 0.05. For each cell line in (B), expression in treated cells is plotted relative to that in untreated cells. The Student’s T-test was used to test for significant differences between untreated and treated cells of the same line. * = P < 0.05, ** = P < 0.01 (C) Immunofluorescence for βIII-tubulin expression in parental and Hmgn2 knockout cell lines plated in the presence or absence of WNT inhibitor and assayed after 48 hrs.

**Figure 5:**
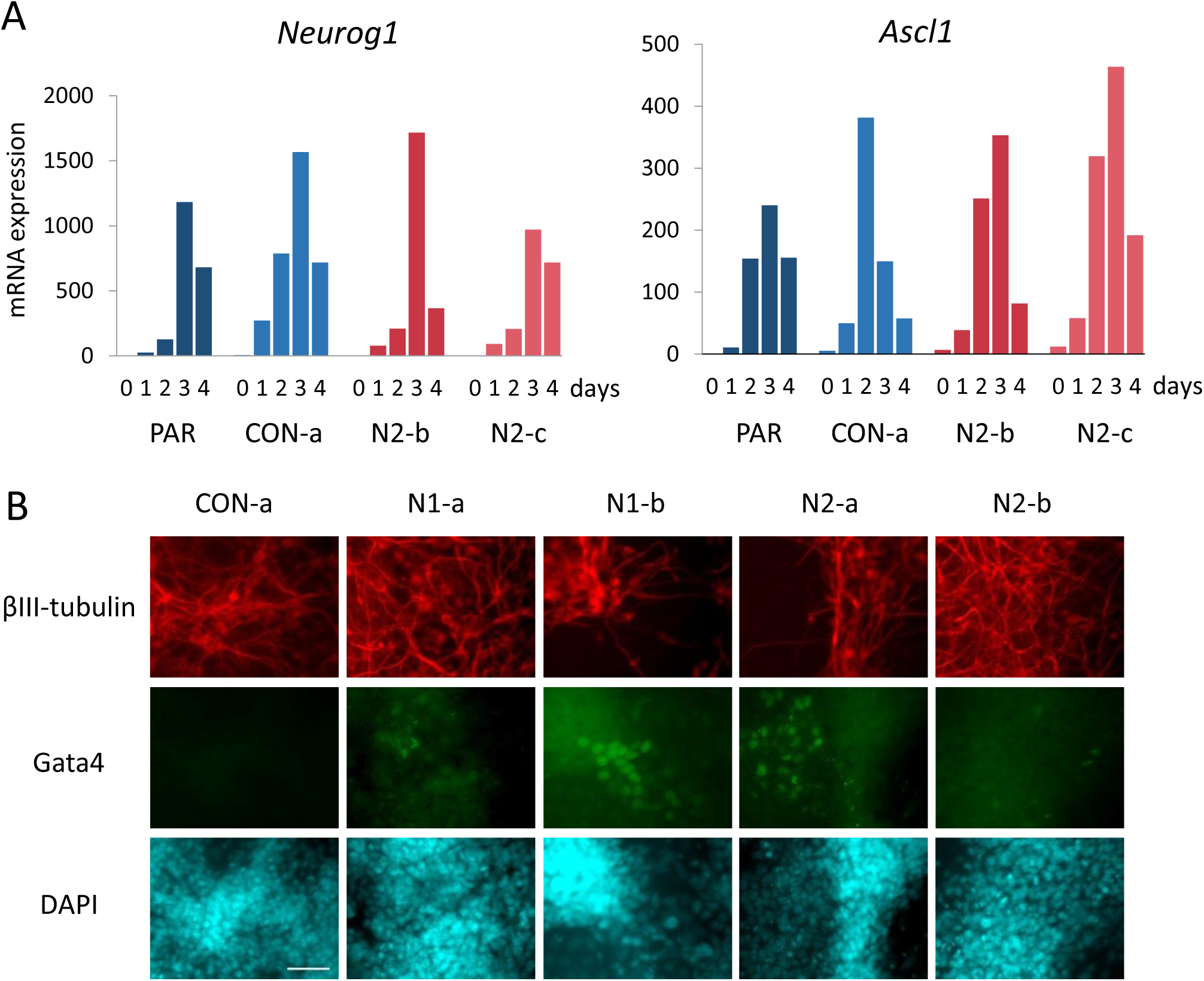
Induced neuronal differentiation is not altered in *Hmgn2* knockout cells. (A) Relative mRNA expression of the *Neurog1* and *Ascl1* genes in parental, CON-a, and N2-b and N2-c cells induced to differentiate along the neuronal lineage. A total of four time points during the protocol were evaluated (1–4 days after induction), together with the undifferentiated cultures (day 0). Expression is plotted relative to that in undifferentiated, untreated parental P19 cells. The data is representative of two independent biological replicates. (C) Immunofluorescence for βIII tubulin, Gata4 and DAPI at day 4 in parental, N1-a, N1-b, N2-b and N2-c cells. Scale bar indicates 50 μm.

*Hmgn1* and *Hmgn2* knockout cells were assayed at several time points in order to assess both differentiation efficiency and timing. Expression of the pro-neural transcription factor *Neurog1* was induced by up to 1500 fold, and *Ascl1* was induced by up to 450 fold during the first four days, but no significant differences were observed between the *Hmgn* knockout cells and the control lines (Figure 5a and supplementary figure S5). Similarly, no significant differences were observed in the timing or level of induction of *Nestin*, nor was the loss of *Nanog* expression during the differentiation process altered significantly (supplementary figure S5).

Immunofluorescence for the neuronal marker βIII tubulin was performed on both *Hmgn1* and *Hmgn2* knockout cells on day 4 after induction, but no significant differences can be observed in the level of marker expression or cell morphology between parental and knockout cells (Figure 5b). Interestingly, a sub-population of Gata4-positive cells was observed in N1-a, N1-b and N2-a cultures, which may indicate initiation of endodermal or mesodermal pathways (Figure 5b).

These results indicate that most of the *Hmgn*-knockout cells respond normally to neuronal induction cues, and are capable of generating neuronal cells with similar timing and efficiency to the parental and control lines.

### Neural stem cells lacking Hmgn1 or Hmgn2 show loss of NSC self-renewal

Although the direct neuronal induction protocol described above is rapid and highly efficient, no glial cells are produced, and the formation of large three-dimensional clusters of cells at later time points makes further analysis more challenging (K.L. West, unpublished). To circumvent these issues, we derived neural stem-like cells from the control and *Hmgn*-knockout P19 cultures (33). NSCs derived from the parental P19 cells can be maintained for many passages, express high levels of Nestin and have a bipolar morphology that is typical of NSCs (PAR-NSC, Figure 6a). Parental P19 NSCs can also be differentiated into mixed cultures of neurons and glia upon removal of the growth factors EGF and FGF (PAR-diff, Figure 6c).

**Figure 6:**
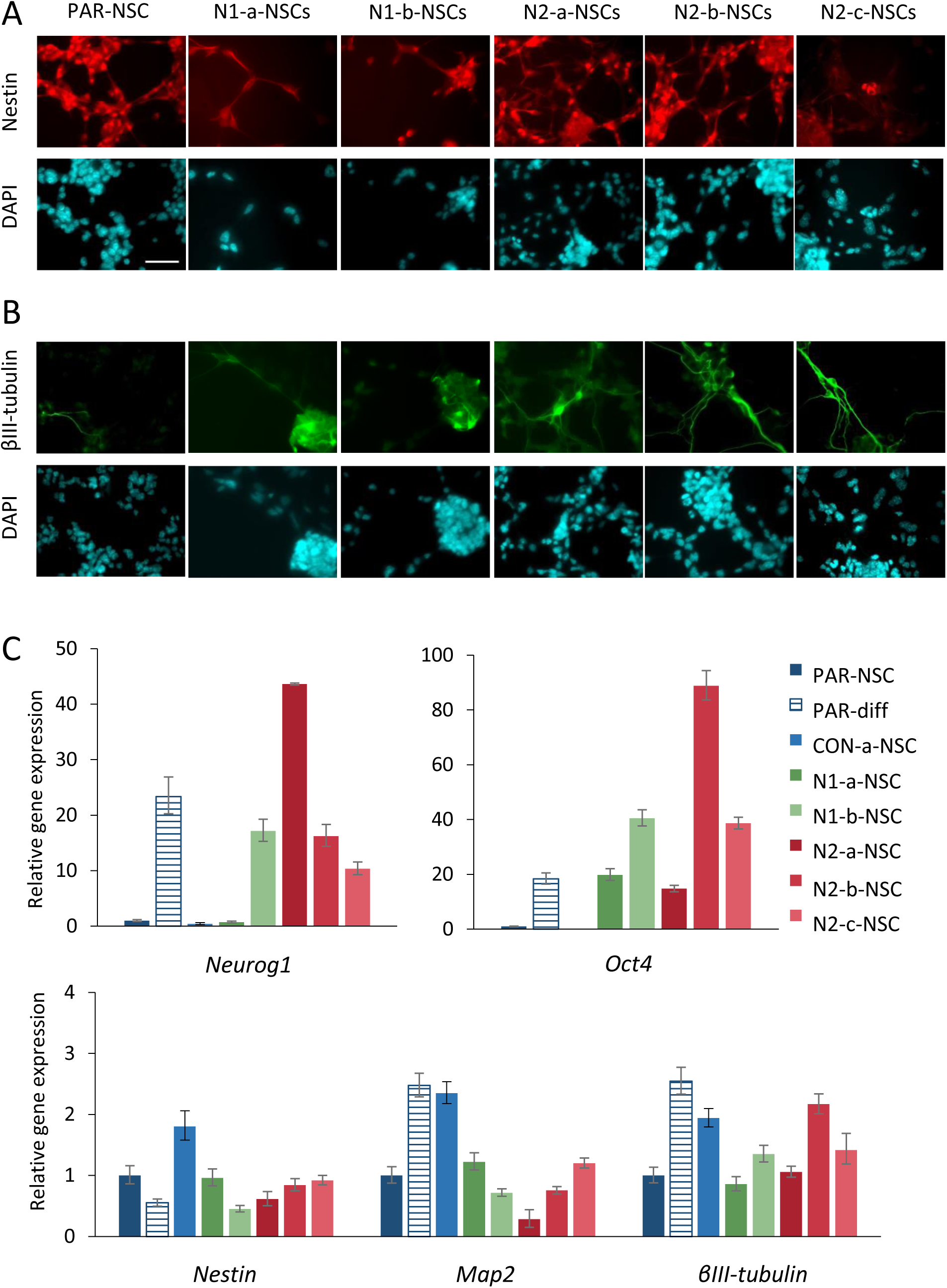
Neural stem cells derived from cells lacking *Hmgn1* or *Hmgn2* show loss of NSC identity. Immunostaining for (A) Nestin and (B) βIII tubulin in NSCs derived from parental P19 cells, *Hmgn1* and *Hmgn2* knockout lines. Scale bar indicates 50 μm. (C) Relative mRNA expression in NSCs derived from parental P19 cells, CON-a cells, and *Hmgn1* and *Hmgn2* knockout lines. Expression in parental NSCs that were induced to differentiate down the neuronal lineage by removal of growth factors is also shown. Expression is normalised to that of *Gpi1*, and is plotted relative to that in parental NSCs. Error bars represent the s.d. from technical qRT-PCR triplicates. Data is representative of two independent NSC derivation experiments.

Immunostaining of *Hmgn*-knockout cultures does not reveal any substantial differences in Nestin expression compared to the parental NSC cultures (Figure 6a). However, higher numbers of βIII-tubulin-positive cells with extended processes are apparent in both *Hmgn1-* and *Hmgn2*-knockout cultures (Figure 6b). At the mRNA level, neither *Nestin* nor *βIII-tubulin* expression is consistently altered in the *Hmgn*-knockout cells, which may indicate that βIII-tubulin protein production is regulated post-transcription in these cells (Figure 6c).

Consistent with the increased numbers of neuronal cells detected by immunofluorescence, mRNA expression of the pro-neural transcription factor *Neurog1* is increased by 10–44 fold in NSCs from N1–1, N2-a, N2-b and N2-c cells compared to the parental NSCs (Figure 6c). This increase is comparable to the 23 fold increase observed when parental NSCs differentiate into neuronal cultures (PAR-NSC versus PAR-diff, Figure 6c). Similarly, *Oct4* expression is increased by 15 – 89 fold in all the *Hmgn1-* and *Hmgn2*-knockout NSCs compared to the control NSCs. This is comparable to the 18 fold increase in *Oct4* expression that is observed when parental NSCs differentiate into neuronal cultures (Figure 6c). However, expression of neuronal marker *Map2* is not consistently altered in *Hmgn* knockout cultures compared to the parental and control NSCs.

These data indicate that Hmgn1 and Hmgn2 play roles in maintaining the self-renewal of NSCs derived from P19 cells. The loss of either variant leads to spontaneous neuronal differentiation, with increased *Neurog1* expression and production of βIII-tubulin-positive cells.

### Loss of Hmgn2 affects the profile of active histone modifications

The data presented earlier shows that loss of Hmgn1 or Hmgn2 in undifferentiated P19 cells affects the expression of the pluripotency gene *Nanog*, the pro-neural genes *Neurog1* and *Ascl1*, and the endodermal/mesodermal markers *Gata4* and *Brachyury*. As Hmgn proteins play several roles in regulating chromatin structure, we first examined publically available ChIP-seq data from mouse ESCs to determine the chromatin structure of these genes (Figure 7c, supplementary figure S6, and data not shown). As expected, the highly expressed *Nanog* and *Oct4* genes have high levels of the active mark H3K4me3 in the vicinity of their TSS, and negligible levels of the repressive mark H3K27me3. Notably, *Neurog1, Ascl1, Gata4* and *Brachyury* have moderate levels of H3K4me3 but also have substantial levels of H3K27me3. This “bivalent” pattern is typical of poised stem cell genes with low levels of expression, which can be upregulated or silenced as the stem cells differentiate (2).

**Figure 7:**
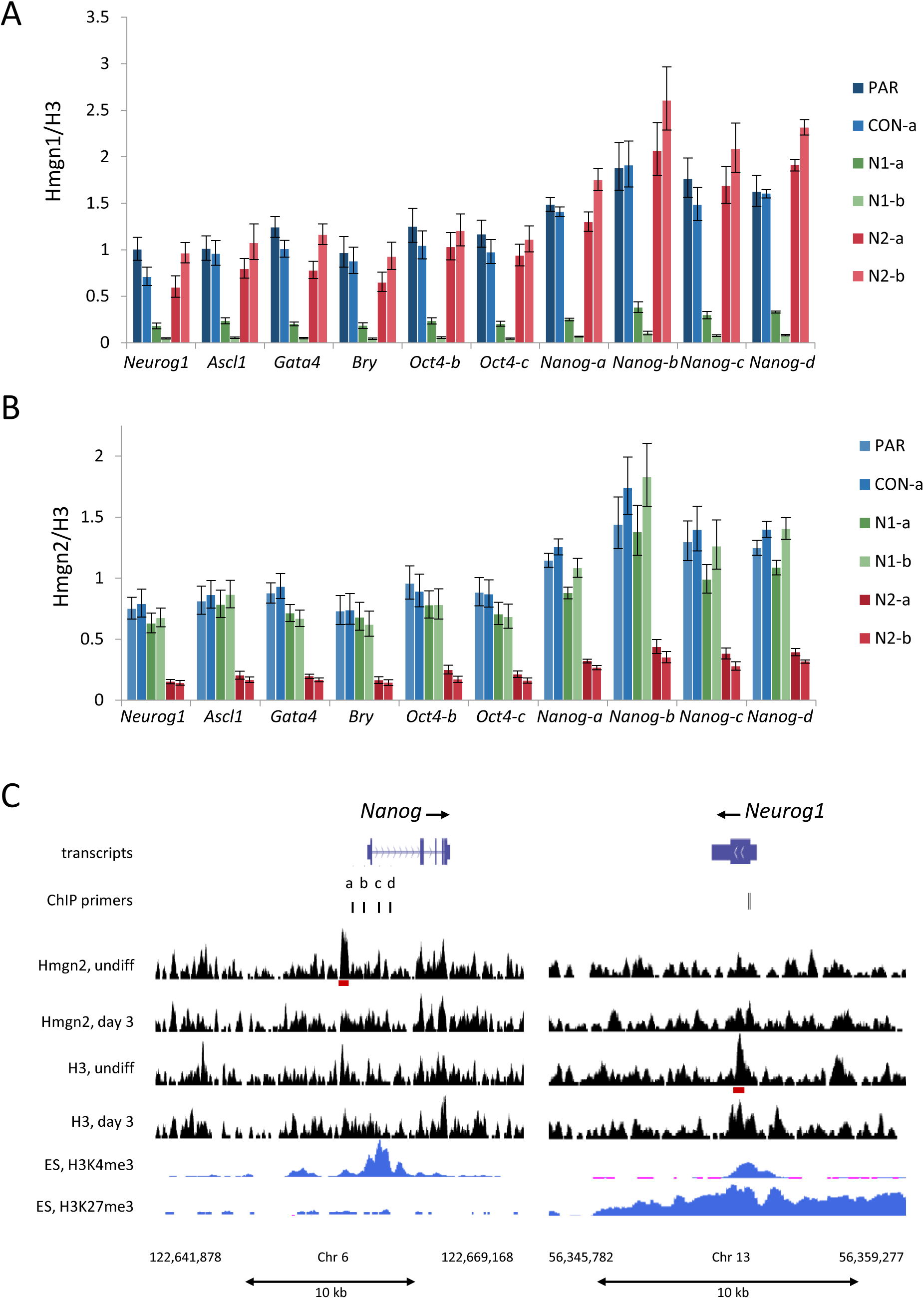
Hmgn1 and Hmgn2 are not highly enriched at active gene promoters. ChIP-PCR assays in parental P19, control CON-a, N1-a, N1-b, N2-a and N2-b cell lines. Enrichment of Hmgn1 (A) or Hmgn2 (B) at each primer set was normalised to the average H3 signal from all primer sets. The graphs present the mean and s.d. of technical qPCR triplicates from the same IP reaction. Data is representative of 2–3 independent biological replicates. (C) ChIP-seq for Hmgn2 and H3 was performed in undifferentiated and day 3 neuronal induced P19 cells. Reads were aligned to the mm9 mouse genome and regions surrounding the *Nanog* and *Neurog1* loci are shown. Peaks revealed by MACS peak calling software for Hmgn2 and H3 are shown as red blocks below the relevant signal track. Positions of the primer sets used for ChIP are indicated. Data for H3K4me3 and H3K27me3 in mouse ES Bruce4 cells was obtained from the UCSC genome browser and was generated by the Ren lab as part of the ENCODE/LICR project (http://genome.ucsc.edu) (55,56). TSSs for the indicated genes were confirmed by comparison with hCAGE data from the FANTOM5 project (57). Y-axis maxima are 0.62 for all H3 and Hmgn2 tracks, 21 for H3K4me3 and 3.5 for H3K27me3.

To investigate whether Hmgn1 or Hmgn2 bind preferentially to the promoters of these genes, chromatin immunoprecipitation followed by qPCR was performed. Between one and four primer sets were designed for each gene locus, using the publically available H3K4me3 data from mouse ES cells as a guide. Single primer sets for *Neurog1, Ascl1*, and *Bry* target the peak of H3K4me3 nearest the TSS of each gene. Binding at the *Oct4* and *Nanog* gene loci was assayed using several primer sets that encompass 2–3 kb surrounding the TSS. The locations of the primers relative to each TSS are illustrated in Figures 7c and S6.

The ChIP data shows that Hmgn1 and Hmgn2 are bound at all the active and poised genes tested. Occupancy varies by less than two fold between different genes, with slightly higher occupancy at the *Nanog* gene locus (Figure 7a and b). In the *Hmgn1* knockout lines, Hmgn1 signals are reduced by 80% and 95% in the N1a and N1b lines, respectively. Conversely, Hmgn2 signals are reduced by 80% on average in the *Hmgn2* knockout lines, N2-a, N2-b and N2-c. These data confirm the specificity of the Hmgn1 and Hmgn2 ChIP assays, and show that these proteins occupy the regulatory regions of both active pluripotency genes and poised/bivalent lineage-specific genes in undifferentiated P19 cells.

In order to investigate whether Hmgn2 binding is particularly enriched at these TSS regions compared to other regions of the genome, ChIP-seq was performed in both undifferentiated P19 cells and P19 cells after three days of induced neuronal differentiation (Figure 7c and supplementary figure S6). The active marks H3K27ac and H3K4me1 were also mapped in order to validate the ChIP assay and illustrate the chromatin landscape of undifferentiated and day 3 neuronal P19 populations (supplementary figure S6). For example, H3K27ac is enriched upstream of the *Oct4* gene in undifferentiated cells, but this is lost in the day 3 neuronal cells when *Oct4* is repressed. Conversely, H3K27ac is absent from the *Ascl1* locus in undifferentiated cells, but increased in the day 3 neuronal cells when the gene is expressed. The H3K27ac and H3K4me1 profiles for undifferentiated P19 cells correspond closely to publically available datasets for undifferentiated mouse ES cells (supplementary figure S6).

The Hmgn2 ChIP-seq binding profiles indicate that Hmgn2 is not highly enriched at active gene promoters, either in undifferentiated or day 3 neuronal cells, but instead is found at similar levels throughout the genome (Figure 7c and supplementary figure S6). In general, the profile of Hmgn2 binding tends to follow that of H3. Peak calling software identified 1,862 Hmgn2 peaks in undifferentiated cells. For example, there is a Hmgn2 peak in the *Nanog* promoter in undifferentiated cells (Figure 7c). However, only 8% of these Hmgn2 peaks overlap with TSSs or putative enhancers.

In order to investigate whether loss of Hmgn2 protein affects histone modifications, ChIP-PCR was performed in undifferentiated parental, N2-b and N2-c cells (Figure 8). At the highly expressed pluripotency genes *Oct4* and *Nanog*, loss of Hmgn2 leads to a loss of active histone marks associated with promoter and enhancer activity. Specifically, the promoter-associated marks H3K9ac and H3K4me3 are reduced at both genes in both Hmgn2 knockout lines, with the reduction in H3K9ac being more substantial (Figure 8a and b). The enhancer-associated marks H3K27ac and H3K122ac are more highly enriched at the *Oct4* locus compared to *Nanog;* both of these modifications are reduced in the *Hmgn2* knockout cells, although the reduction in H3K27ac levels is greater (Figure 8d and f). These reductions in active histone modifications do not appear to have a strong influence on expression of the genes, however, as *Oct4* expression is unchanged in the N2-b and N2-c lines, whereas Nanog levels are decreased in N2-c cells. *Oct4* and *Nanog* have negligible levels of the repressive mark H3K27me3 in parental cells, and this is not altered in the *Hmgn2*-knockout lines (Figure 8c).

**Figure 8:**
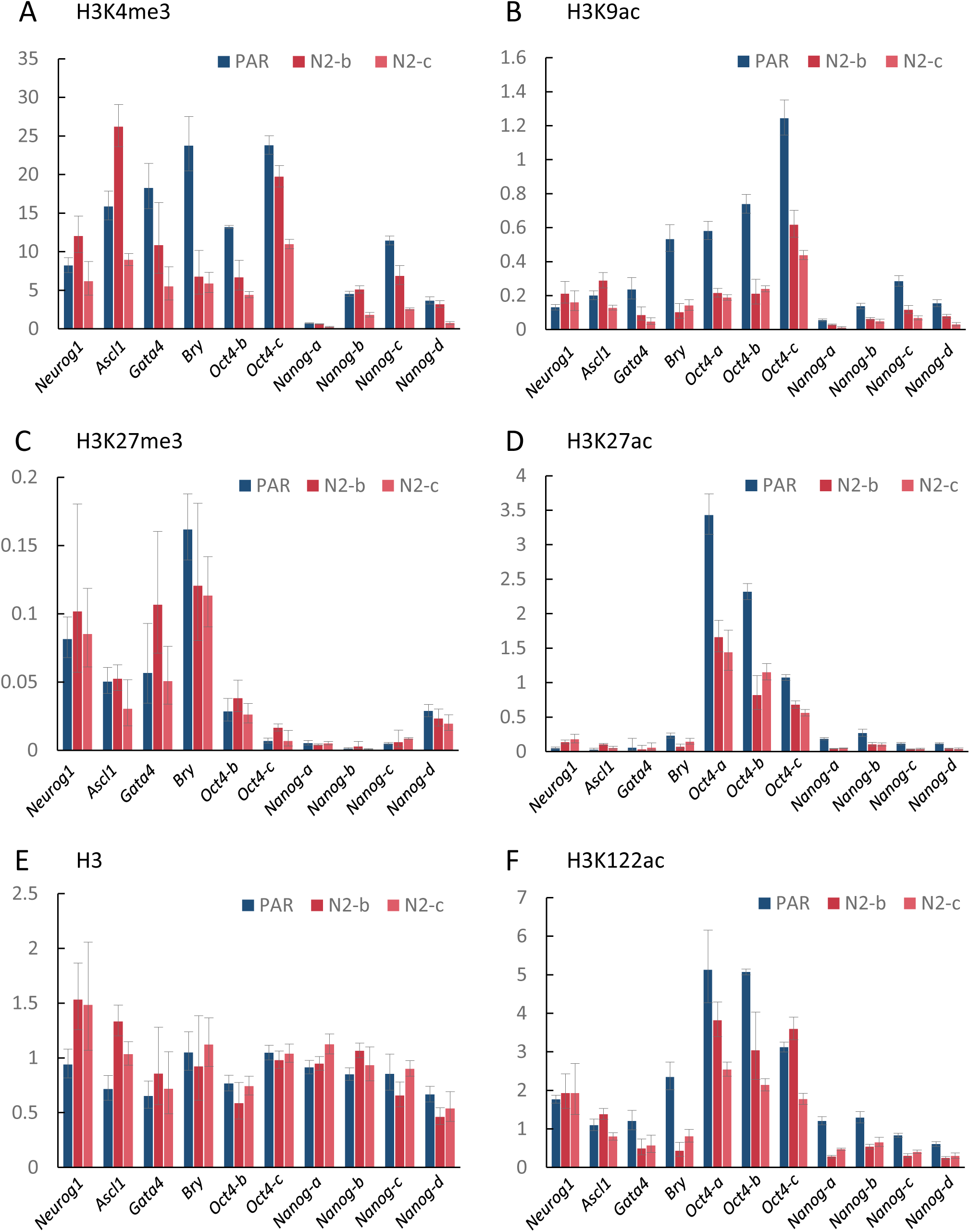
Hmgn2-Ko cells show a reduction in active histone H3 modifications at bivalent and active gene loci. ChIP-PCR assays in parental P19, N2-b and N2-c cell lines for H3K4me3 (A) and H3K9ac (B), H3K27me3 (C), H3K27ac (D), H3 (E) and H3K122ac (F). The enrichment of each modification was normalised to the average H3 signal from all primer sets. The graphs present the mean and s.d. of technical triplicates from the same IP reaction. Data is representative of 2–3 independent biological replicates.

The bivalent endodermal/mesodermal marker genes, *Gata4* and *Brachyury*, lose the active promoter marks H3K4me3 and H3K9ac in *Hmgn2*-knockout cells, but retain the repressive H3K27me3 mark (Figure 8a, b, c). These changes indicate a switch from bivalency to a more repressive chromatin state. This does not appear to impact *Gata4* expression, which is unchanged in the N2-b and N2-c cells. However, the reduction in active marks is greater at the *Brachyury* locus, which is consistent with the observed decrease in *Brachyury* expression in these lines (Figure 3d).

Conversely, pro-neural genes whose expression increases in Hmgn2 null cells do not lose their bivalent signature. H3K9ac is unchanged at either *Neurog1* or *Ascl1*, and H3K4me3 at *Ascl1* is increased in N2-b cells and decreased in N2-c cells. No changes in H3K27me3 were observed either locus (Figure 8a, b, c).

In order to extend the observations made at individual genomic locations, the global levels of several active histone modifications were investigated in *Hmgn1* and *Hmgn2* knockout cells (supplementary figure S7). Consistent with the ChIP data, global levels of H3K4me3 are reduced by 64% in N2-b cells compared to control CON-a cells, but this reduction is not observed in the other *Hmgn* knockout lines tested. Both the N2-a and N2-b knockout lines show a significant reduction in global H3K9ac levels, consistent with the ChIP data, but this is not observed in the *Hmgn1* knockout lines. The greatest reduction was observed with H3K27ac, which is significantly reduced in both *Hmgn1* (N1-a) and *Hmgn2* (N2-b) knockout cells (supplementary figure S7).

In summary, it is clear that loss of Hmgn2 can lead to reductions in the levels of active promoter and enhancer histone modifications at specific gene loci. The highly expressed pluripotency genes *Oct4* and *Nanog* show a reduction in active marks at promoters and enhancers. The poised endodermal/mesodermal genes *Gata4* and *Brachyury* switch from bivalency to a more repressive chromatin configuration, whereas the poised pro-neural genes *Neurog1* and *Ascl1* retain their bivalent state. Some, but not all, of these changes are reflected in the global levels of the modifications, with H3K27ac being most strongly reduced in both *Hmgn1* and *Hmgn2* knockout lines.

## Discussion

### Hmgn proteins regulate the self-renewal of pluripotent and multipotent embryonal carcinoma cells

Hmgn proteins are highly expressed during development and in differentiating cells, but are downregulated in mature tissues (15,22). In particular, high Hmgn expression has been observed in the neural stem/progenitor cells of the sub ventricular zone of the mouse brain, and the transit-amplifying progenitor cells of hair follicle (3,22,25,26). A role for Hmgn proteins in regulating the self-renewal or differentiation of stem/progenitor cells is supported by the observation that downregulation of Hmgn expression is important for the differentiation of limb bud mesenchymal cells into chondrocytes *in vitro* (15,24).

Here, we have studied the role of Hmgn1 and Hmgn2 in pluripotent embryonal carcinoma cells, and in neural stem cells derived from them. With both *Hmgn1* and *Hmgn2* knockout EC cells, we observe increases in spontaneous neuronal differentiation, which are accompanied by the loss of some pluripotency markers and increases in the expression of pro-neural transcription factors.

Bright field microscopy and immunostaining revealed that Hmgn-knockout cells are morphologically heterogeneous and are less likely to grow in tight colonies, resembling the differentiated cells in ESC cultures (37). A high proportion of *Hmgn*-knockout cells are devoid of the pluripotency markers Ssea1 and Nanog, as revealed by immunofluorescence. Nanog is one of several pluripotency-related transcription factors that are downregulated in the transition from naïve pluripotency to the primed state for differentiation (29,30). However, Nanog is known to be heterogeneously expressed in ESC and P19 cultures, and the low-Nanog cells show higher lineage-specific gene expression (38). Although the low-Nanog cells are more prone to differentiate, they have not abandoned pluripotency as they indefinitely self-renew and can generate daughter cells with higher Nanog levels (38). Therefore, it seems unlikely that the increased number of differentiated cells in the Hmgn knockout cultures is due solely to a reduction in Nanog expression.

Expression of the transcription factors Oct4 and Sox2 is not altered by the loss of Hmgn1 or Hmgn2. Unlike Nanog, Oct4 and Sox2 are retained during the early stages of differentiation where they interact with different lineage-specific transcription factors to influence fate commitment (40). It is possible that some of the Oct4-positive cells in the Hmgn knockout cultures have abandoned pluripotency and initiated differentiation programs.

Spontaneous differentiation of *Hmgn*-knockout cells down the neural lineage is evidenced by βIII-tubulin immunostaining, although some Gata4-positive cells are also observed in some of the lines. Consistent with this, gene expression analyses show increased *Neurog1* and *Ascl1* expression in *Hmgn1* and *Hmgn2* knockout lines. The predominance of neuronal cells over other lineages is consistent with the default model of neural differentiation, which proposes that pluripotent embryonic stem cells commit to neural fates unless instructed otherwise (49). For example, ESCs start losing their identity upon withdrawal of the factors that maintain them in a pluripotent state, and they differentiate mostly into neural lineages (33). Neurog1 and Ascl1 are powerful pro-neural regulators whose overexpression can drive the neuronal differentiation of P19 cells and ES cells (50,51). It is possible that these two transcription factors are key contributors to the spontaneous neuronal differentiation observed in the *Hmgn*-knockout cells.

Our observations contrast with previous work performed in mouse ES cells derived from *Hmgn1^−/−^* and *Hmgn1^−/−^n2^−/−^* mice, where ES cell pluripotency and self-renewal did not appear to be altered (3,12). It is unclear whether the differences are related to the transformed nature of the EC cells, or due to the growth media used, which may permit or restrict spontaneous differentiation depending on whether cells are cultured in the presence of serum or 2i inhibitors. It is also worth considering the different developmental stages from which ESCs and P19 cells were derived. ESCs capture the naïve state of pluripotency from pre-implantation embryos, whereas epiblast stem cells represent a pluripotency state primed for differentiation (29–31). If this is valid for transformed cells, P19 cells that were derived from post-implantation embryos can immediately commit to differentiation, whereas ESCs first need to exit the naïve pluripotency stage so that they may respond to inductive cues (30). Therefore, Hmgn proteins may not be relevant in the preservation of the naïve pluripotency; rather, these chromatin architectural proteins may be important for maintaining the primed cells unresponsive to misplaced inductive differentiation cues.

To investigate whether the loss of self-renewal that we observed in the undifferentiated cultures was specific to pluripotent stem cells, we generated multipotent neural stem/progenitor cells from the control and *Hmgn* knockout cell lines. Notably, the gene expression profiles of the neural stem cell cultures were altered in *Hmgn1* and *Hmgn2* knockout cells, with substantial increases in *Oct4* and *Neurog1* expression, and increased levels of βIII-tubulin – positive neuronal cells. Thus, the loss of control of self-renewal is not specific to pluripotent P19 cells, but is also apparent in P19-NSCs lacking *Hmgn1* or *Hmgn2*. It has been previously demonstrated that *Hmgn1^−^*^/^*^−^*mice have fewer Nestin-positive cells in the sub-ventricular region of the brain (3), which is consistent with the hypothesis that self-renewal is compromised in neural stem/progenitor cells lacking Hmgn1 or Hmgn2. Our results complement a previous study that demonstrated a role for Hmgn proteins in regulating the differentiation of neural stem/progenitor cells (NPCs) (26). Overexpression of Hmgn proteins in NPCs, both *in vivo* and *in vitro*, was shown to promote astrocyte differentiation and inhibit neurogenesis, whereas knocking down *Hmgn* expression had the converse effect (26).

### Neuronal signalling pathways are not altered in Hmgn knockout EC cells

The timing of differentiation is regulated by transcriptional switches that are driven by intrinsic or extrinsic signals. Three major pathways that regulate neuronal differentiation are Notch, FGF and WNT. Inhibitors were used in order to investigate whether changes in these signalling pathways are driving the differentiation of Hmgn knockout cells. Known target genes of these pathways, *Axin, Fgf4* and *Hes5*, are not significantly altered in the Hmgn knockout cells, and treatment with the inhibitors has similar effects on these target genes in both the control and the knockout cells, indicating that these pathways are operating normally in the Hmgn knockout cells.

Notch signalling promotes self-renewal and survival of NS cells in early stages of development, whereas in later stages it favours glial versus neuronal fates (45). Inhibition of neuronal differentiation by the Notch pathway is mediated by its targets, Hes1 and Hes5, which suppress the expression of pro-neuronal transcription factors such as Neurog1, Neurog2 and *Ascl1* (48). However, Notch is not relevant for pluripotency (52), which explains the observation that *Nanog* expression is not influenced by iNotch in any of the cell lines. Inhibition of Notch signalling led to a small increase in *Neurog1* expression of up to 3 fold in both the control and the *Hmgn2* knockout lines, indicating that Notch does not play a major role in preventing neuronal differentiation of P19 cells.

FGF and WNT pathways establish a cross-talk to pattern the neural plate and influence cell fate (53). FGF signalling is used to derive and maintain the NS cells, but also to induce neuronal differentiation *in vitro* (46). Fgf4 stimulates FGF signalling in an autocrine manner, allowing embryonic pluripotent cells to commit to differentiation (54). In agreement, *Nanog* is upregulated in all P19 cell lines after inhibition of FGF signalling, suggesting that at least some of the cultured cells can revert to a more naïve pluripotency state under FGF signalling inhibition. The fact that interfering with FGF signalling has a comparable effect in all the cell lines, including higher pro-neural gene expression, suggest that the spontaneous neural differentiation of Hmgn2-knockout cells is not explained by changes in FGF signalling.

WNT signalling has also been reported to have dual roles during neuronal differentiation; in the expansion phase it promotes self-renewal of NS cells, but in the neurogenic phase it acts as an instructive pathway for neuronal differentiation by targeting the pro-neural transcription factor Neurog1 (47). WNT signalling also does not seem to be responsible for the neural differentiation of P19 cells after the loss of *Hmgn2*, as iWnt has little effect on pro-neural gene expression. In conclusion, it appears that none of the tested signalling pathways trigger differentiation of P19 cells after the loss of Hmgn2, as interfering with them does not re-establish lineage-specific gene expression to parental levels. These observations suggest that the loss of a major Hmgn variant is permissive for the initiation of differentiation following an intrinsic and cell-autonomous program.

### Hmgn proteins modulate the epigenetic landscape of EC cells

The binding of Hmgn proteins to nucleosomes is known to be transient and dynamic (17). Here, ChIP-seq shows Hmgn2 is bound throughout the non-repetitive genome, and with no highly enriched peaks at transcription start sites or enhancers in pluripotent P19 cells and early neuronal cultures. Its binding profile is broadly similar to that of the core histone H3. Previous studies have indicated that Hmgn proteins are enriched at DHSs in some cell types but not in others, although it is not clear whether the different antibodies used have contributed to the apparent differences in binding profiles (3,9,12,15,16,18–20). It is conceivable that the accessibilities of specific Hmgn epitopes may vary, depending on the Hmgn protein conformation or its binding partners. We then performed more detailed chromatin analysis at three classes of genes expressed in pluripotent stem cells – pluripotency associated genes (*Nanog, Oct4*), bivalent endodermal/mesodermal lineage genes (*Brachyury, Gata4*) and bivalent pro-neural genes (*Neurog1, Ascl1*). The data confirm that both Hmgn1 and Hmgn2 are bound at the promoter regions of all three gene classes of gene.

Chromatin immunoprecipitation assays show that loss of Hmgn2 leads to a reduction in the active histone modifications at the promoters and enhancers of these target genes. However, the extent of the changes and the impact on gene expression differs between each gene class. The levels of H3K4me3, H3K9ac, H3K27ac and H3K122ac are reduced at the pluripotency associated genes *Nanog* and Oct4; this corresponds to a reduction in *Nanog* expression, although *Oct4* levels are unchanged. H3K4me3 and H3K9ac are reduced at the poised genes *Gata4* and *Brachyury*, such that their bivalent chromatin state is switched to a more H3K27me3-dominant repressive chromatin configuration. However, this only appears to reduce *Brachyury* expression, as *Gata4 expression* is unchanged in these cell lines. Conversely, the pro-neural genes *Neurog1* and *Ascl1*, whose expression is increased in the *Hmgn* knockout cells, show no changes in histone modifications and retain their bivalent status. Genome-wide studies would be required to investigate whether these trends are also found at other highly expressed and poised genes. However, it is clear that the loss of Hmgn2 significantly impacts the epigenetic landscape of P19 cells, leading to a reduction in active marks at specific gene loci. The data supports a role for Hmgn2 in maintaining the open chromatin conformation at gene regulatory elements in embryonic stem cells, thus supporting the chromatin plasticity that is required for self-renewal and pluripotency (1).

Several previous studies have shown that Hmgn proteins directly influence the levels of histone H3 modifications. Specifically, Hmgn1, Hmgn2 and Hmgn3a can stimulate the ability of PCAF to acetylate nucleosomal H3K14, with Hmgn2 showing the greatest effect (6,8,9). This activity is dependent on the C-terminal regulatory domain, and kinetic and structural data reveal that interaction of the Hmgn protein with the H3 N-terminal tail may make it a better substrate for histone acetylase activity (8,11). Conversely, Hmgn proteins have been shown to inhibit phosphorylation of H3 at serine 10 and serine 28 (6,7). Further investigation is needed to determine whether Hmgn proteins can also influence the deposition or removal of H3K4 methylation.

Deng *et al*. have previously shown that loss of both Hmgn1 and Hmgn2 leads to increased levels of the repressive mark H3K27me3 around the *Olig1* and *Olig2* gene loci in mES cells (12). This is mediated by increased H1 binding in the absence of Hmgn proteins, leading to increased recruitment of the H3K27 methyltransferase, EZH2. We do not observe increased H3K27me3 levels in P19 cells lacking a single Hmgn variant, possibly because levels of the remaining Hmgn variant are sufficient to prevent increased H1 binding.

We conclude that the loss of Hmgn2 leads to a widespread reduction in the levels of active modifications at histone H3. However, the epigenetic changes at individual gene loci do not always match the changes in gene expression observed in these cells. Further work is required to determine whether other features such as nucleosome positioning, histone variant deposition or transcription factor recruitment are altered in the *Hmgn* knockout cells and could affect the transcriptional profile and cellular phenotype (10,13,14).

The results of the present work demonstrate that the loss of Hmgn1 or Hmgn2 compromises the ability of embryonal carcinoma cells, and the NSCs derived from them, to self-renew accurately and maintain pluri/multipotency. As a consequence, a proportion of the cells exit pluri/multipotency and initiate differentiation programs. We show that the loss of Hmgn proteins disrupts the active modification landscape in pluripotent stem cells, which impacts the transcription of both pluripotency genes and bivalently-marked differentiation factor genes. Our data is consistent with a growing consensus that Hmgn proteins do not act as specific regulators of gene expression, but fine-tune a cell’s transcriptional profile by influencing the landscape of histone modifications and chromatin structure (3,12,19). We propose that Hmgn proteins contribute to chromatin plasticity in pluripotent stem cells, facilitating open chromatin states that are essential for maintaining self-renewal and pluripotency.

## Availability of data

The ChIP-seq datasets supporting the conclusions of this article are available in the NCBI GEO repository under accession number GSE118031.

**Supplementary Data are available at NAR online.**

## Acknowledgements

We thank Narendra Kumar and Tony McBryan for assistance with the bioinformatic analysis, and we thank Faika Laz Banti and Martin Šrámek for additional lab work. Many thanks to all members of the KLW and AGW labs for their assistance, advice and support.

